# Capsid-mediated chromatin state of the AAV vector genome controls host species range

**DOI:** 10.1101/2022.10.06.511169

**Authors:** Adriana Gonzalez-Sandoval, Shinnosuke Tsuji, Feijie Zhang, King L. Hung, Howard Y. Chang, Katja Pekrun, Mark A. Kay

## Abstract

Recombinant adeno-associated viral vectors (rAAVs) are among the most commonly used vehicles for in vivo based gene therapies. However, it is hard to predict which AAV capsid will provide the most robust expression in human subjects due to the observed discordance in vector-mediated transduction between species. We used a primate specific capsid, AAV-LK03, and demonstrated that the limitation of this capsid towards transduction of mouse cells was unrelated to cell entry and nuclear transport but rather due to depleted histone H3 chemical modifications related to active transcription, namely H3K4me3 and H3K27ac, on the vector DNA itself. A single-amino acid insertion into the AAV-LK03 capsid enabled efficient transduction and the accumulation of active histone marks on the vector chromatin in mouse without compromising transduction efficiency in human cells. Our study suggests that the capsid protein itself is involved in determining the chromatin status of the vector genome, most likely during the process of uncoating. Programming viral chromatin states by capsid design may enable facile DNA transduction between vector and host species.

## Main Text

Adeno-associated virus (AAV) is a small single-stranded DNA (ssDNA) virus of the Parvoviridae family. While the discovery of a single capsid serotype and mechanisms related to basic AAV biology were described as early as 1960 ^1,2^, in recent times recombinant AAV vectors have become the most popular vehicles for in vivo based gene therapy. Since the capsid sequence determines host tropism, a lot of research over the past decades has focused on creating vectors with novel properties to be effective in human gene therapy applications. Apart from isolating new AAV variants from natural reservoirs the capsid sequence has been modified either by rational design or directed evolution to develop capsids with specific properties. One of the major limitations has been the discordance in transduction efficiencies among species. Several studies have shown that rAAVs with capsids that can be used with high efficiency in preclinical mouse models are commonly found to be less efficient in non-human primate studies and/or human clinical trials^3^. In some cases, AAV capsids have been efficient in primates but not in rodents^4^, thus making the use of a surrogate capsid for testing in preclinical mouse studies necessary. The unpredictability of serotype specificity adds time, cost, and uncertainty to the research process of developing effective gene-based therapeutics^5^.

We decided to characterize the AAV transduction process in primate and rodent species, to better understand the primate selectivity observed with some AAV capsids. We focused on the AAV-LK03 capsid because of its current use in clinical trials^6^. AAV-LK03 is an engineered capsid originating from a capsid-shuffled library which had been selected in a xenograft humanized liver mouse model^4^. AAV-LK03-mediated gene transfer results in poor transgene expression in mice but performs robustly in primates including humans^6^. In our study we compared different stages of the AAV-LK03 transduction process between both species. We found that the low transduction efficiency of AAV-LK03 in mouse cells was not due to major differences in nuclear accumulation of vector genomes but was rather correlated with a lack of histone modifications related to active transcription. Our study identifies epigenetic regulation as part of the species selectivity of AAV capsids. Our hypothesis was supported by the observation that addition of a single amino acid in the AAV-LK03 capsid, which restored transduction efficiency in murine cells, was associated with the accumulation of active-related epigenetic marks without marked changes in the amount of nuclear AAV genomes. Our results support previous findings that defining transduction efficiency requires both the number of vector copies in the target cell and transgene expression. This study reveals the importance of developing AAV capsids that enable formation of active chromatin in the cargo vector genome.

### AAV-LK03 derived genomes internalize but do not express in mouse cells

To identify the mechanism of preferential activity of AAV-LK03 in primate cells, we assayed different steps of the transduction process *in vitro* and *in vivo* (Fig. 1). We used AAV-DJ (a chimeric capsid isolated from a capsid shuffled library, selected on human hepatoma cells) as a control capsid, as it is known to transduce both primate and rodent cells^7^. Huh7 and Hepa1-6 hepatoma cell lines were utilized as representative cells of human and mouse origin, respectively. A constitutive luciferase expression cassette (CAG-Luciferase) was packaged with AAV-LK03 and AAV-DJ capsids and the two cell lines were transduced with the resulting rAAV vectors. As expected, AAV-LK03 resulted in 100x lower luciferase activity in the mouse Hepa1-6 cells as compared to the human Huh7 cells, while AAV-DJ showed similar luciferase activity in both cell lines (Fig. 1 a). Luciferase activity was shown to correlate with relative luciferase mRNA levels (Fig.1 b). However, when quantifying the vector copy number in whole cell lysates and fractionated nuclear lysates (Fig. 1 c, d), internalization of the LK03 packaged vector in the nucleus was only ~3-fold reduced in the mouse cells as compared to the human cells. On the other hand, similar levels of AAV-DJ delivered vector DNA was found in both cell lines. We confirmed that those observations were not unique to these particular cell lines or expression cassette, as similar results were obtained using vectors with different promoters and transgenes in various human and mouse cell lines (Extended Data Fig 1 a-c). Similar results were obtained *in vivo*. The livers of mice, which had been subjected to systemic delivery of the CAG-Luciferase vector packaged with AAV-LK03 vs AAV-DJ capsids showed only a 3-fold reduction in nuclear vector copy number while transgene expression (mRNA and protein) was >100x reduced (Fig. 1 f-i).

**Fig. 1:**
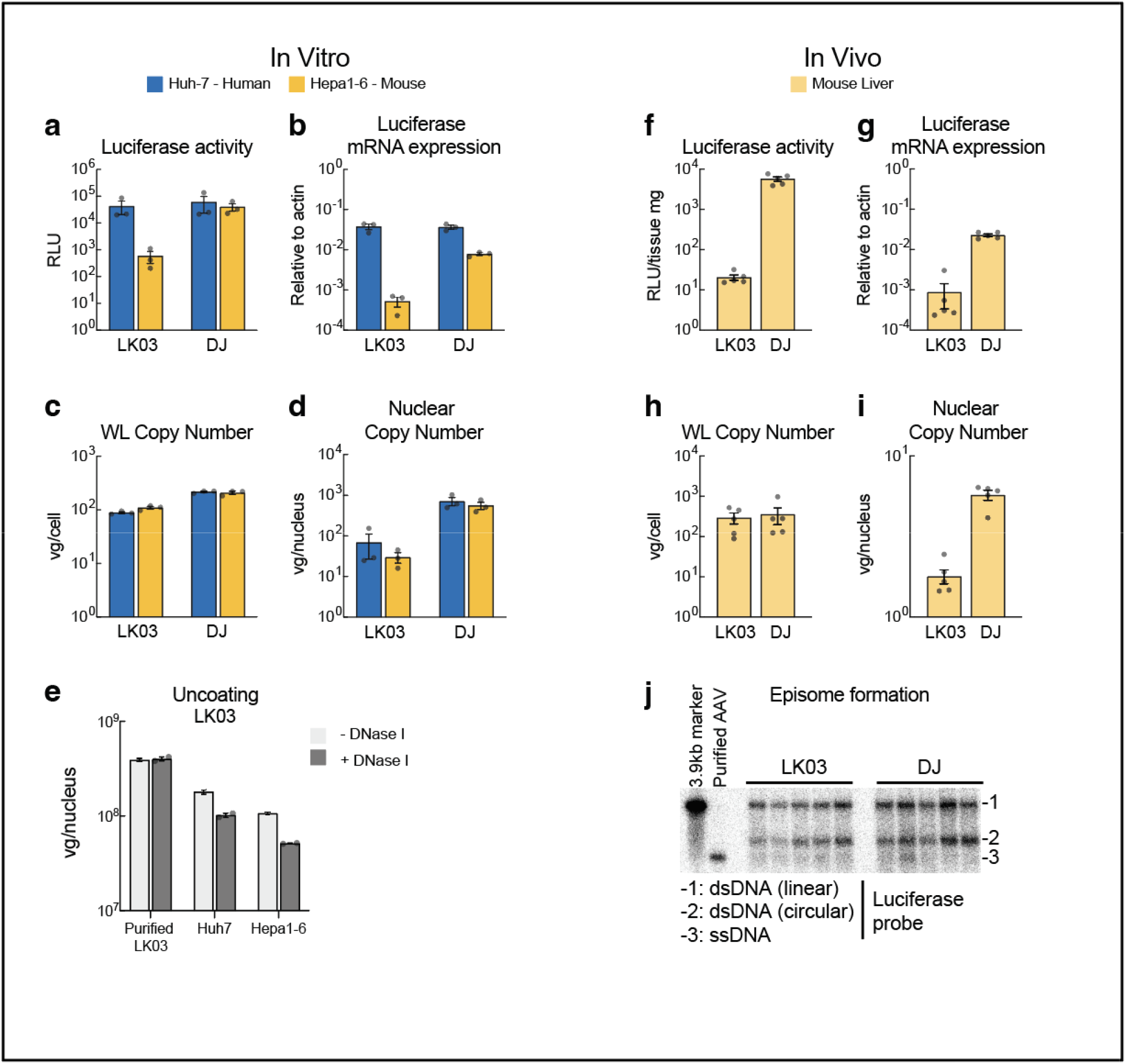
Characterization of AAV-LK03 transduction in human and mouse hepatoma cells. **(a-e)** *In vitro* assays using cell lines 48h post transduction, **(f-j)** *in vivo* assays using liver tissue 3 day post rAAV delivery (i.v. tail vein). **(a, f)** Luciferase activity quantification. **(b, g)** Relative quantification of Luciferase mRNA by qRT-PCR. **(c, h)** Quantification of cellular vector genome by qPCR in whole cell lysate (WL) or **(d, i)** in nuclei. **(e)** Determination of the uncoating efficiency of AAV-LK03 by quantification of vector copies in DNase I treated vs untreated nuclear extracts. **(j)** Southern blot analysis in liver tissue probing for the luciferase gene.

As part of the transduction process, before the transgene can be expressed, the ssDNA AAV genome needs be released from the AAV capsid in a process known as uncoating. The ssDNA is then converted into an episome, primarily as circular double-stranded DNA (dsDNA)^8–10^. We compared the uncoating efficiency of AAV-LK03 between species by treating transduced cells with DNase I prior to DNA extraction to degrade the uncoated vector genomes^11^ (Fig. 1e). We found that a similar proportion of vector genomes was encapsidated and thus protected from degradation in the +DNase I condition in both species. Analysis of transduced mouse liver (Fig. 1j) revealed that the quantity and conformation of AAV-LK03 derived episomes were comparable to those derived from the DJ capsid, suggesting that the steps of uncoating and episome formation were not the main reasons for poor AAV-LK03 mediated transgene expression in mouse liver. In addition, when we used a self-complementary (sc) rAAV vector to circumvent the need for double strand formation, a similar level of discordance in transduction efficiency (transgene expression) between species was observed (Extended Data Fig. 1d). Thus, episome formation was unlikely to be the main restriction factor for the observed species specificity of AAV-LK03.

### AAV-LK03 delivered genomes lack active histone marks

Given the comparable number of episomes derived from both AAV-LK03 and AAV-DJ *in vivo*, we reasoned that the ~3-fold decrease in nuclear internalization cannot be the primary reason for the >100-fold difference in mRNA or luciferase activity. We hypothesized that differences in chromatin structure and/or histone modification between the genomes delivered by different capsids might influence the silencing or activation of episomal vectors. We performed Cut&Tag^12^ to quantify the genome-wide landscape of histone modifications, including those related to active transcription (H3K4me3 and H3K27ac) and those related to transcriptional repression (H3K9me3 and H3K27me3)^13^. AAV-LK03 delivered genomes were depleted of H3K4me3 in the gene body region, with only a minor enrichment around the ITRs in mouse cells (Fig. 2a). As a control, enrichment of H3K4me3 within the albumin (*Alb*) gene was similar in all samples transduced with either capsid, regardless of the species transduced (Extended Data Fig. 2a). Looking at all histone modifications and conditions (Fig. 2b), plotted as coverage (which considers enrichment, read length and genomic size region), we found that AAV-LK03 delivered genomes were depleted for both histone modifications linked to active transcription (H3K4me3 and H3K27ac) in mouse cells, while the repressive histone modifications (H3K9me3 and H3K27me3) were poorly enriched on AAV delivered genomes in both species irrespective of the capsid used. This suggests that the reduced transcription from AAV-LK03 does not stem from the accumulation of repressive-related histone modifications, but rather from the lack of active histone modifications.

**Fig. 2:**
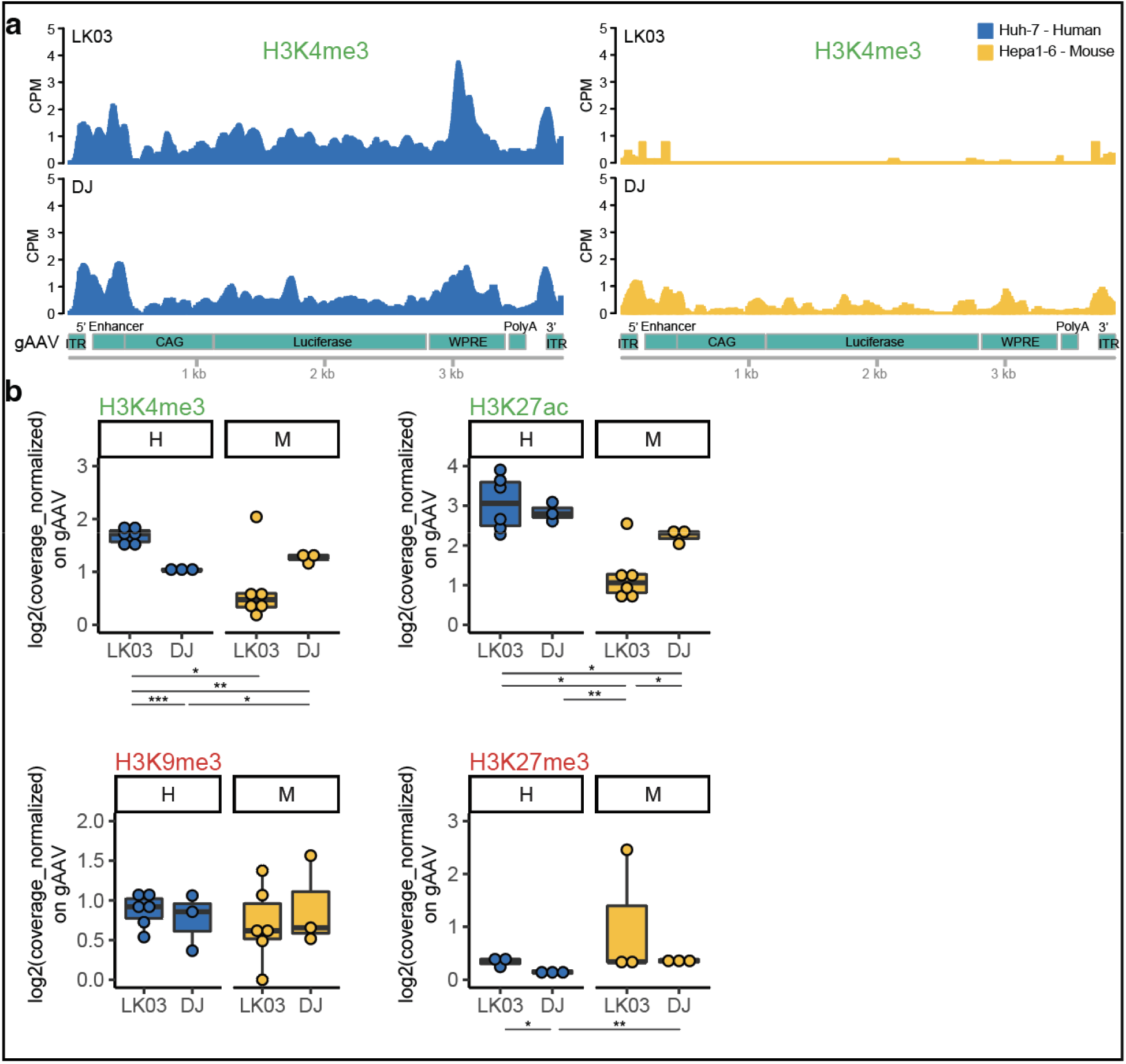
Species-specific histone modifications in rAAV genomes delivered with the AAV-LK03 capsid. **(a)** H3K4me3 normalized (CPM) signal on AAV genome obtained from the Cut&Tag assay and next generation sequencing for Huh7 human and Hepa 1-6 mouse cell line transduced with AAV-LK03 and AAV-DJ, as indicated. **(b)** Cut&Tag boxplots of normalized coverage on the AAV genome, for histone epigenetic modifications - H3K4me3 and H3K27a (active transcription), H3K9me3 and H3K27me3 (inactive transcription) for Huh7 human and Hepa 1-6 mouse cell lines transduced with AAV-LK03 and AAV-DJ. Statistic p-value * <0.05, ** <0.01, *** <0.001. H=human, M=mouse.

To interrogate if the low enrichment of histone modifications in mouse cells transduced with AAV-LK03 was caused by a lack of core histones, we performed Cut&Tag targeting core histones H2A, H3 and H4 (Extended Data Fig 2). AAV-LK03-delivered genomes showed comparable levels of core histones in all experimental conditions, suggesting proper nucleosome assembly on AAV-LK03 delivered genomes in mouse cells.

### Single point mutant of AAV-LK03 capsid allows delivered genomes to contain active-related epigenetic marks

Capsid sequence alignment revealed loss of a glycine or threonine residue from the highly variable region around serine 264 to serine 267 (S264-S267) in capsids with preferential activity in primates over mice^14^ (Extended Data Fig. 3a). AAVLK03-265insT^15^ is an example of improved mouse liver tropism of AAV-LK03 by insertion of a single amino acid in this variable region, similar to what has been described for the closely related AAV3B^14^. However, this particular insertion leads to a considerable increase in nuclear vector copies in mouse cells (Extended Data Fig 3 b) and, therefore, may obscure mechanisms of capsid-specific species tropism other than the difference in DNA copies. Thus, we created AAV-AM by inserting a glycine residue at position 266. According to a predictive structure model of this new serotype compared to the parental AAV-LK03 capsid (AAV3B) crystal structure (Fig. 3 a), most of the capsid structure remains unchanged, except for an extension of the VR-I loop which allows for closer proximity of the adjacent alanine and serine side chains to the histidine side chain which is six amino acids downstream. Delivery of the luciferase expression cassette using the AAV-AM capsid restored luciferase expression in mouse cells to levels similar to those observed in human cells. Expression levels were also comparable to those achieved with AAV-LK03 in human cells (Fig. 3b). The enhanced expression was not the result of increased vector copy numbers (Fig. 3 c), which were nearly identical between AAV-AM and AAV-LK03 in mouse cells. However, in human cells we observed ~10-fold higher vector genome uptake using AAV-AM compared to AAV-LK03, suggesting that AAV-AM influences nuclear internalization in a species-specific manner. We also observed enrichment of active histone marks on vector genomes delivered by AAV-AM, both for mouse as well as for human cells (Fig. 3 d). The repressive histone modifications H3K9me3 and H3K27me3 were comparable to other serotypes (Extended Data Fig. 3 b,d). Core histones were also present in vector genomes delivered by AAV-AM (Extended Data Fig 3 c,d).

**Fig 3:**
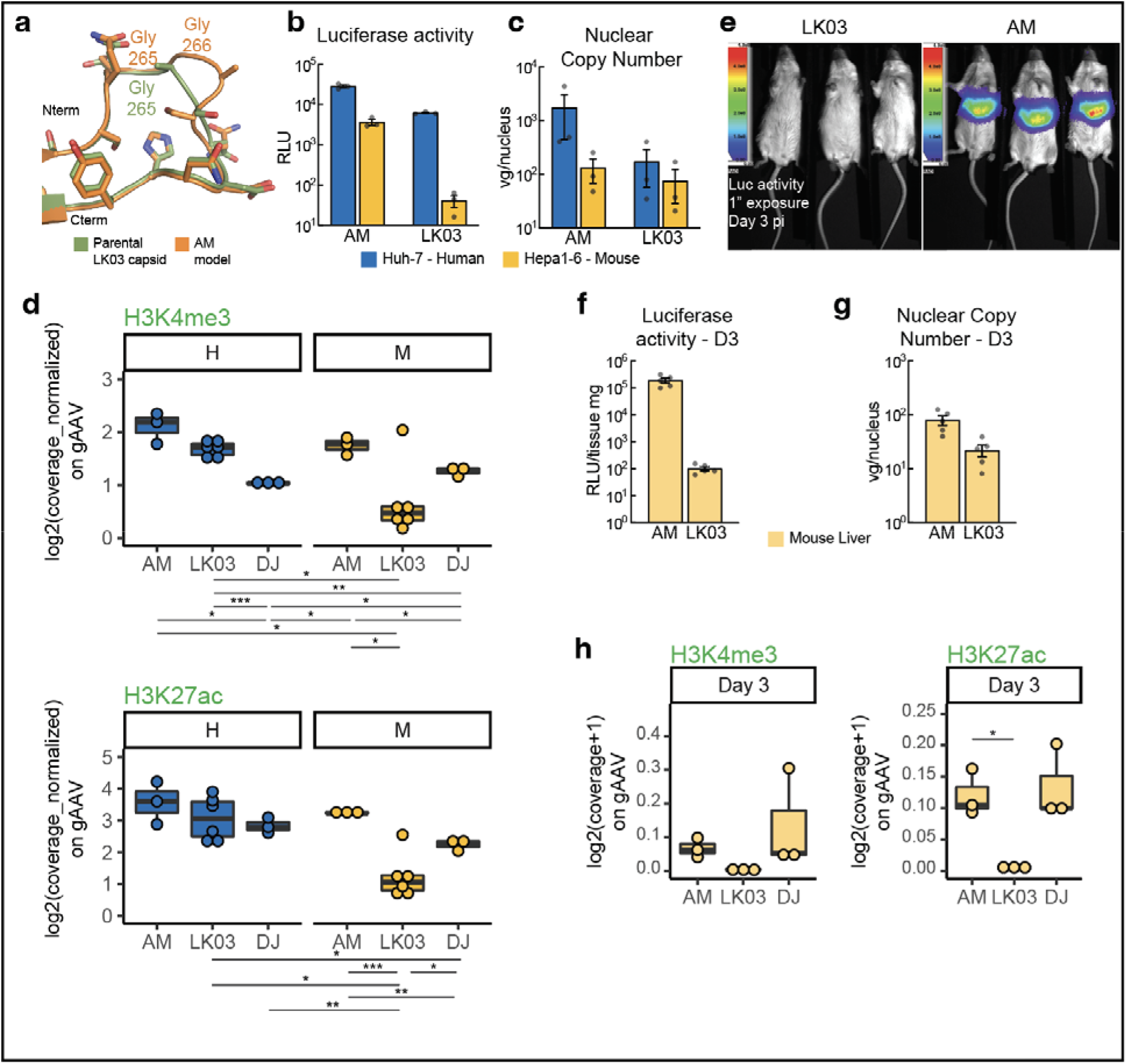
A single glycine amino acid insertion in AAV-LK03 (AAV-AM) promotes transduction in mice and active histone marking. **(a)** Homology model built using SwissModel server using the 3kie crystal structure as template (Parental AAV-LK03 capsid - AAV3B). **(b)** *In vitro* 48h transduction luciferase activity assay. **(c)** Nuclear copy number quantification in Huh7 human and Hepa 1-6 mouse cells transduced with AAV-LK03 and AAV-AM by qPCR. **(d)** Cut&Tag boxplots of normalized coverage on AAV genome of active-related histone marks in vitro. **(e)** *In vivo* luciferase activity assay 3 days post intravenous injection of AAV-LK03 and AAV-AM. **(f)** Liver tissue luciferase assay. **(g)** Nuclear copy number qPCR quantification 3 days post intravenous vector injection. **(h)** Cut&Tag boxplots of normalized coverage on AAV genome of active-related histone marks in vivo. Statistic p-value * <0.05, ** <0.01, *** <0.001. H=human, M=mouse.

Importantly, high levels of luciferase activity were detected in livers of mice that had been intravenously injected with AAV-AM (Fig. 3 e, f). When compared to mice that had been injected with vector packaged with AAV-LK03 capsid nuclear internalization was only ~3-fold increased (Fig. 3 g). We also performed Cut&Tag on mouse liver for the three different capsids. Enrichment of active-related histone marks was observed for AAV-AM and AAV-DJ, but not for AAV-LK03 (Fig. 3 h). As a reference, the *Albumin* gene enrichment of H3K4me3 epigenetic mark was similar in all mouse liver samples transduced with either capsid (Extended Data Fig. 4 a). Inactive-related histone modifications (Extended Data Fig 4 b), and core histones (Extended Data Fig 4 c) enrichments were also comparable between AAV-AM and AAV-DJ, while AAV-LK03 delivered genomes exhibited lower levels of enrichment.

**Fig 4:**
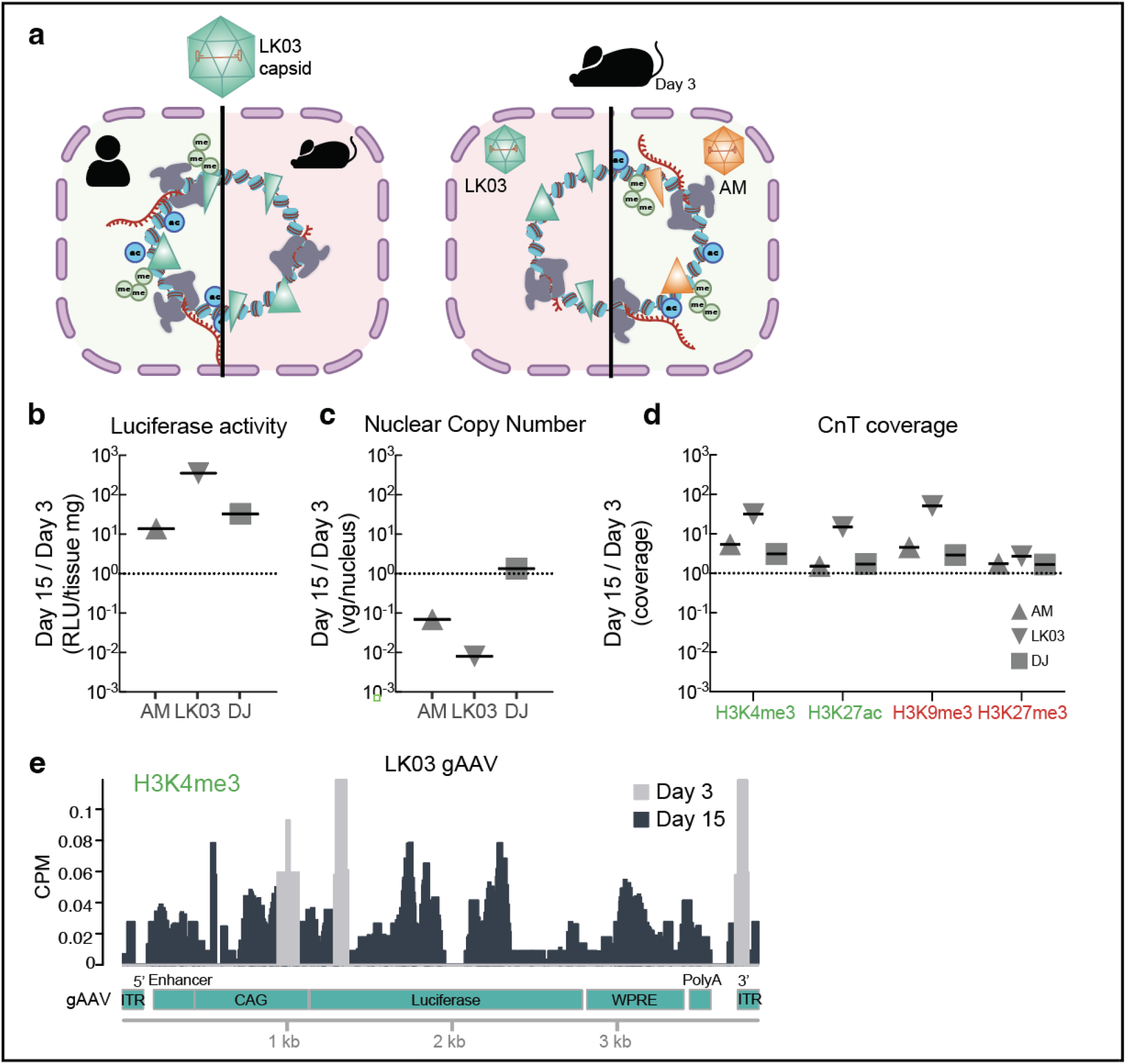
Species selectivity of AAV-LK03 mediated transgene expression involves the formation of active-related histone epigenetic marks. **(a)** Schematic representation of results. AAV-LK03 derived genomes were enriched for active-related epigenetic marks in human, but not mouse cells. After adding a single amino acid to the AAV-LK03 capsid (AAV-AM), transduction wa efficient in mouse cells and liver. Active-related epigenetic marks were enriched in AAV-AM transduced mice. **(b-d)** Ratio comparisons of in vivo assays on mouse liver tissue at Day 15 v Day 3 post intravenous vector injection: **(b)** Luciferase activity, **(c)** Nuclear copy number RT-qPCR, **(d)** Cut&Tag coverage of histone epigenetic marks as indicated. **(e)** Comparison of Cut&Tag H3K4me3 normalized (CPM) signal on the AAV vector genome at different days post injection as indicated.

Our data strongly supports the hypothesis that the failure of AAV-LK03 to efficiently transduce rodent derived cells and tissues is related to a lack of transcription-permissive histone modifications associated with the episomes. In contrast, the AAV-AM capsid, which contains a single amino acid insertion as compared to the AAV-LK03 capsid, was able to cause the vector genome to accumulate activating epigenetic marks allowing transgene expression in mouse cells as well as in human cells (Fig. 4 a). These results lead us to propose that the capsid interacts with host cell proteins during the uncoating process and helps to establish the epigenetic state of the vector episome. We also determined that these interactions differ between species for AAV-LK03 but not for AAV-AM.

### Remnant episomes derived from LK03 transduction in mouse liver contain active-related histone modifications

We sought to examine the stability of AAV episomes and their epigenetic state over time. Since episomal DNA is lost during cell division in cell culture studies, we examined the enrichment of histone modifications and core histones in mouse livers harvested 15 days post injection with CAG-FLuc vectors packaged with the three different capsids. Comparing day 15 vs day 3 harvested livers, luciferase expression increased for all three AAV vectors (Fig. 4 b, Day 15 Luciferase activity in Extended Data Fig 5 a), yet the gAAV nuclear copy number was reduced in a capsid-specific manner: AAV-AM was reduced 10-fold and AAV-LK03 100-fold during the 12-day interval (Fig. 4 c, Day 15 Nuclear copy in Extended Data Fig 5 c).

Nevertheless, we observed an increase in coverage of histone modifications on vector genomes independent of capsid (Fig. 4 d), higher enrichment of active-related histone modifications than inactive-related and found the largest differences in enrichment for AAV-LK03 derived genomes. Core histones did not vary as much as the histone modifications (Extended Data Fig 5c). Interestingly, AAV-LK03 delivered genomes at day 15 post injection contained histone modifications across the entire gAAV, in contrast to the pattern observed at day 3 (Fig. 4 e and Extended Data Fig. 5 d-f). Although the enrichment level on day 15 was not higher overall than on day 3, it was unexpected because the vector copy number was reduced 100-fold in mice transduced with AAV-LK03 as compared to those injected with AAV-DJ. We suggest that AAV-LK03 derived genomes that persist overtime require a favorable deposition of histones and their modifications.

## Discussion

The discordance observed by us and others regarding AAV-transduction efficiency between species has contributed to some of the difficulties in gene-based therapeutics. The use of AAV-LK03 requires the use of surrogate capsids in preclinical rodent testing, which is not optimal. Our results further exemplify the importance of defining transduction and should include the measurement of transgene expression either by mRNA and/or protein production, as measuring vector genomes alone can be misleading. We had previously shown that DNA plasmids delivered into mouse liver cells via hydrodynamic transfection become chromatinized^16^ so it is not surprising that chromatinization of rAAV vectors occurs once they uncoat in the nucleus and are converted to episomes.

However, it is remarkable that the AM capsid, a derivative of the primate specific AAV-LK03 harboring a single amino acid insertion, resulted in delivering similar numbers of episomes into mouse cells but result in > 100x improvement in transgene expression. Using Cut&Tag, an assay to uncover epigenetic marks in a high-throughput format, we were able to establish that the differential expression patterns were related to the relative enhancement in transcriptionally permissive histone marks.

These results show that the capsid proteins play a role in the transduction efficiency of vector genomes. Capsid proteins being important for transduction efficiency have been slowly gaining some spotlight in other contexts as well, where various capsids have differential effects on expression from various promoters used in the central nervous system^17^. Others have reported a point mutant of the capsid AAV2 that can internalize into cells and uncoat at similar levels as wildtype AAV2 but fails to result in transgene expression ^18^.

Our results underscore the importance of histone association and epigenetic regulation of vector genomes, resulting in retention in the nucleus and active transcription due to accumulation of activating histone marks. We propose that the nuclear uncoating process and histone association with the vector genome is a crucial step in the transduction mechanism and is dependent on the capsid sequence.

Our data also implicates the type of epigenetic marks placed on the vector genomes is dependent on how the capsid proteins interact with chromatin modifiers, which might be different between species and/or various cell types. Identification of these host factors will enable us to understand more details about how AAV genomes can become chromatinization with active epigenetic marks, which will lead to more rational design/screening of novel capsids with both clinical/preclinical utility. One example consistent with general epigenetic regulation is the silencing of AAV2 derived genomes by NP220 and the HUSH complex^19^, indicating that perhaps the active- or inactive-related modifications might be capsid dependent. In a humanized liver model comparisons between human xenograft vs mouse resident hepatocytes showed a block in human transduction efficiency of AAV-DJ occurring post-uncoating^20^. In contrast to the mouse hepatocytes a significant proportion of the human hepatocytes harbored AAV-positive nuclei despite a lack of transgene mRNA expression^18^. It is possible that higher mammals have evolved more sophisticated processes for episomal epigenetic regulation and perhaps there are polymorphic variant genetic factors within primates that may in part explain the wide range of responses in AAV-mediated gene transfer between individuals. Taken together, the studies mentioned above suggest potential differences in capsid-mediated epigenetic regulation between species. Further investigation will be needed to understand the detailed mechanism by which the capsid proteins influence the deposition of various modified histones and perhaps other chromatin modifiers on the vector genome.

## Acknowledgments

We thank W. Greenleaf and G. Marinov for helpful discussions. A.G-S. is grateful to S. Herrera, A. Gatto for help and support in bioinformatic analysis. A.G-S is thankful to I. Deshpande for help in modelling of AAV-AM capsid. This project was also supported by a NIH Shared Instrumentation Grant (S10-OD010580) from the National Center for Research Resources (NCRR) with significant contribution from Stanford’s Beckman Center as well as the Stanford Small Animal Imaging Facility. The authors wish to acknowledge the Stanford Genomics Facility for performing high-throughput sequencing, as well as CMP and SCG facilities.

## Author contributions

A.G-S., S.T. and M.A.K. designed the research. S.T, carried out initial experiments. F.Z. carried out mouse injections and liver harvesting. K.P participated in the in vivo luciferase assays and provided several reagents for AAV production. K. L. H. developed the initial computational analysis, which was later adjusted by A.G-S. A.G-S., S.T., K.L.H., analyzed data. H.C. provided discussion. A.G-S, K.P., and M.A.K. wrote the manuscript.

## Methods

### Vector plasmids

The AAV vectors containing ITR sequences used in this study are based on AAV type 2 backbone. Most experiments use rAAV vectors expressing Firefly Luciferase (FLuc) under control of a CAG promoter (pAAV-CAG-FLuc). The cloning of pAAV-CAG-FLuc (Addgene Cat# 83281) plasmid has been described previously^21^.

An experiment uses rAAV vectors expressing FLuc under EF1a (core) promoter (pAAV-EF1a-FLuc). The pAAV-EF1a-FLuc plasmid was generated by replacing the TdRed sequence in pAAV-EF1a-TdRed (unpublished, see below) with the FLuc gene with the regulatory WPRE region from pAAV-CAG-FLuc (Addgene Cat# 83281). Restriction enzymes BamHI-HF and SalI-HF were used to release the 3.4 kb vector band from pAAV-EF1a-TdRed as well as the 2.3kb FLuc-WPRE sequence from pAAV-CAG-FLuc. Both fragments were ligated with Hi-T4 ligase and transformed into One Shot Stbl3 competent E.coli. The unpublished pAAV-EF1a-TdRed vector was generated by replacing the RSV promoter region in plasmid pAAV-RHB^22^ with the first 212 nucleotides of the EF1a promoter from plasmid pAAV-EF1a-FLuc-WPRE-HGHpA (Addgene Cat# 87951) and the hAAT transgene with the TdRed sequence obtained from plasmid pAAV-CAG-tdTomato (Addgene Cat# 59462).

An experiment uses rAAV vectors expressing FLuc under CMV promoter (pAAV-CMV-FLuc). The cloning of pAAV-CMV-FLuc plasmid was prepared by swapping the CAG promoter from pAAV-CAG-FLuc (Addgene Cat# 83281) with the CMV promoter from pCMV6-Entry (Origene Cat# PS100001). BamHI-HF and NdeI restriction enzymes were used for both plasmids, the 5.6kb and 0.4kb fragments were isolated, ligated with Hi-T4 ligase and transformed in One Shot Stbl3 competent *E.coli*.

An experiment uses rAAV vectors expressing GFP (pAAV-CAG-GFP (Addgene Cat# 37825)) or TdTomato (pAAV-CAG-TdTomato (Addgene Cat# 59462)) under CAG promoter.

An experiment uses self-complementary rAAV vectors expressing RLuc under CAG promoter from pscAAV-CAG-RLuc (Addgene Cat# 83280).

The packaging vector AAV-LK03 (AAV2 rep LK03 cap) has been described previously^4^. AAV-LK03insT and AAV-AM were created by inserting a Threonine or Glycine respectively immediately downstream of the Serine at position 264 of the AAV-LK03 capsid using the QuikChange in vitro mutagenesis kit (Agilent).

### AAV production

rAAV vectors were produced as previously described using a triple transfection protocol with PEI 25K^23^, followed by purification by CsCl gradient^23^ or using an AAVpro Purification kit (all serotypes; Takara), following the manufacturer’s instruction. Purified rAAVs were stored in aliquots at −80⍰°C until use. rAAV genomes were extracted and purified using a QIAamp MinElute Virus Spin kit (Qiagen) and titered by qPCR with serial dilutions of a plasmid standard. qPCR was performed with 2uL of corresponding material, in duplicate using Apex qPCR Green Master Mix (Genesee Scientific) and a CFX384 Touch Real-Time PCR Detection System (Bio-Rad) using the following cycling conditions: 95⍰°C for 15⍰min, 45 cycles of 95⍰°C for 10⍰s, 60⍰°C for 10⍰s and 72⍰°C for 10⍰s and one cycle of 95⍰°C for 10⍰s and 65⍰°C for 1⍰min and 65–97⍰°C (5⍰°C s–1). The sequence information for primers used in qPCR is shown in Supplementary Table 1.

### Cell culture and transduction

Huh7 cells were purchased from JCRB, 293T and Hepa1-6, SNU-499, PLC, C3A, Hepa-1c1c7, BNL, AML12 cells were purchased from ATCC. Huh7, Hepa 1-6, Hepa-1c1c7, BNL lines were cultured in DMEM with 10% fetal bovine serum (FBS), 2⍰mM L-glutamine and 2mM sodium pyruvate; additional 22mM of HEPES buffer is added to 293T cells. SNU-499 were cultured in RPMI-1640 medium. PLC, C3A were culture in EMEM with 10% FBS. AML12 were cultured in DMEM-F-12 with 10% FBS. Cells were seeded and transduced after 24h with a MOI (multiplicity of infection) of 1e4 vg/cell, and incubate for further 48h for most of experiments, unless indicated for 24h.

### Animals and transduction

All animal work was performed in accordance with the guidelines for animal care at Stanford University. BALB/c scid mice (Strain #:001803) were purchased from Jackson Laboratory. We used 6 week-old juvenile male mice. rAAV delivery was accomplished by tail vain injection with 3e11vg per mouse. Mice were housed at 18.3–23.9⍰°C, with 40–60% humidity on a 12-h light/12-h dark cycle. Food and water were given ad libitum.

At the end of each experiment, mice were anesthetized with isoflurane and perfused transcardially with PBS, and liver tissues were quickly collected and cut into several pieces. The tissues for mRNA extractions were immediately submerged in RNAlater solution (Sigma-Aldrich) and stored at 4⍰°C until use. For luficerase, gDNA or Cut&Tag assays, tissues were snap-frozen in liquid nitrogen and stored at −80⍰°C until use.

### Firefly luciferase assays

Luciferase assays in vitro were performed using the ONE-Glo Luciferase Assay System (Promega) following the manufacturer’s instruction. Briefly, at indicated time points after rAAV transduction, 50uL of the reconstituted substrate was added to the cells grown in 96-well plates and incubated for 10⍰min with gentle shaking. Luminescent activity was measured using a plate reader and Tecan i-control Microplate Reader Software.

*In vivo* luciferase imaging of mice was performed by intra peritoneal injection of 150 μg per g body weight D-Luciferin (Biosynth, catalog L-8220) and ventral luciferase readings using an Ami Imaging System.

Luciferase assay of liver tissue was performed measuring tissue weight (ranging from 50-300mg), homogenizing tissue in RINO 1.5-ml Screw-Cap tubes filled with stainless steel beads (Next Advance Inc, NC0542451) and 200uL of 1x Passive Lysis buffer (Promega) using a bead homogenizer (Next Advance Bullet Blender Storm - BBY24M) at speed 8 for 3 minutes. Cell debris was pelleted by centrifugation at 10’000 rpm for 10min at 4C and supernatants were collected as liver extract. 5uL of liver extract were added to 100uL of ONE-Glo Luciferase reconstituted substrate and incubated for 10⍰min with gentle shaking. Luminescent activity was measured using a plate reader and Tecan i-control Microplate Reader Software. Values were normalized by tissue weight. Three technical replicas were performed for each liver tissue.

### RNA extraction, cDNA preparation and RT-qPCR

Total RNA was extracted using a RNeasy micro plus kit (Qiagen) according to the manufacturer’s protocol with on-column DNase treatment for 30 minutes.

For *in vitro* experiments, cultured cells transduced with rAAVs were collected by trypsinization and washed with PBS. At least 1e6 cells were used for RNA extraction.

Liver tissue samples stabilized in RNAlater solution (~100⍰mg) were homogenized in RINO 1.5-ml Screw-Cap tubes filled with stainless steel beads (Next Advance Inc, NC0542451) and 800⍰μl of RLT buffer (including β-mercaptoethanol) using a bead homogenizer (Next Advance Bullet Blender Storm - BBY24M). 600uL of lysate were used for total RNA extraction. cDNA was synthesized from 50–100⍰ng of total RNA using a High-Capacity RNA-to-cDNA kit (Life Technologies) according to the manufacturer’s instructions.

qPCR was performed with 2uL of 1:5 diluted cDNA, in duplicate using Apex qPCR Green Master Mix (Genesee Scientific) and a CFX384 Touch Real-Time PCR Detection System (Bio-Rad) using the following cycling conditions: 95⍰°C for 15⍰min, 45 cycles of 95⍰°C for 10⍰s, 60⍰°C for 10⍰s and 72⍰°C for 10⍰s and one cycle of 95⍰°C for 10⍰s and 65⍰°C for 1⍰min and 65–97⍰°C (5⍰°C s– 1). Targets of interest were normalized against ActinB (human or mouse) as Delta-Delta Ct calculations. All sequence information of primers is listed in Supplementary Table 1.

### gDNA extraction

Total gDNA was extracted using a QIAamp DNA Mini kit (Qiagen) in cultured cells and DNeasy Blood & Tissue kit (Qiagen) in liver tissue, according to the manufacturer’s protocols with addition of RNase A treatment.

Cultured cells transduced with rAAVs were collected by trypsinization and washed with PBS. At least 1e6 cells were used for total gDNA extraction.

Snap-frozen liver tissue (~100⍰mg) was homogenized in RINO 1.5-ml Screw-Cap tubes filled with stainless steel beads (Next Advance Inc, NC0542451) and 200uL of 1x Passive Lysis buffer (Promega) using a bead homogenizer (Next Advance Bullet Blender Storm - BBY24M) at speed 8 for 3 minutes. Centrifuged at 10’000 rpm for 10min at 4C, collecting supernatant as liver extract. 100uL of liver extract were used for total gDNA extraction.

### Nuclear gDNA extraction

Cultured cells transduced with rAAVs were collected by trypsinization and washed with PBS. Nuclei were isolated with NE-PER™ Nuclear and Cytoplasmic Extraction Reagents (ThermoFisher Scientific). Briefly, the cell pellet was resuspended in CER I buffer + Halt Protease Inhibitor cocktail (ThermoFisher Scientific) in a volume according to pellet size as per manufacturer’s instructions. The suspension was vortexed, incubated on ice for 10 min and vortexed again. Then, CER II was added according to the manufacturer’s instructions, the lysate was vortexed, incubated for 1 min on ice and vortexed again. Nuclei were pelleted by 3 min centrifugation at 13’000 rpm at 4C, washed with 100uL of CER I buffer, and x3 times with 500uL of cold PBS spinning for 1 min at 13’000 rpm. After adding 200uL of ALT Buffer + 20uL Proteinase K, the nuclear pellet was sonicated for 5 min and genomic DNA was extracted using the QIAamp DNA Mini kit (Qiagen) according to the manufacturer’s instructions. The additional RNase A treatment was included in the protocol.

Snap-frozen liver tissue (~100⍰mg) was homogenized in RINO 1.5-ml Screw-Cap tubes filled with stainless steel beads (Next Advance Inc, NC0542451) and 1ml of CER I buffer using a bead homogenizer (Next Advance Bullet Blender Storm - BBY24M) at speed 8 for 3 minutes. The extract was transferred into a fresh tube, incubated on ice for 10 min, and vortexed. Subsequently 55uL of CER II were added, the mixture was vortexed, incubated for 1 min on ice and vortexed again. Nuclei were pelleted by 3 min spin at 13’000 rpm at 4C, the nuclear pellet was washed with 100uL of CER I buffer, and 3 times with 500uL of cold PBS, and spun for 1 min at 13’000 rpm. Finally, 200uL of ALT Buffer + 20uL Proteinase K were added, the nuclear pellet was sonicated for 5 min and nuclear gDNA was extracted following the protocol of the DNeasy Blood & Tissue kit (Qiagen) with addition of RNase A treatment.

### Vector copy number qPCR

Nuclear or whole lysate DNA was used from cultured cells or liver tissues. qPCR was performed with 2uL of eluted DNA, in duplicate using Apex qPCR Green Master Mix (Genesee Scientific) and a CFX384 Touch Real-Time PCR Detection System (Bio-Rad) using the following cycling conditions: 95⍰°C for 15⍰min, 45 cycles of 95⍰°C for 10⍰s, 60⍰°C for 10⍰s and 72⍰°C for 10⍰s and one cycle of 95⍰°C for 10⍰s and 65⍰°C for 1⍰min and 65–97⍰°C (5⍰°C s–1). Standard curves for each primer set were generated using serially diluted plasmid (for gAAV) or Human and Mouse Genomic DNA (for host) (Promega G1521 and G3091, respectively) and used for quantification. CFX Maestro Software was used for data analysis. All sequence information of primers is listed in Supplementary Table 1.

### Uncoating - DNase I treatment

Cultured cells transduced with rAAVs were collected by trypsinization and washed with PBS. Nuclei were extracted as described in the previous section (Nuclear gDNA extraction), then the nuclear pellets were resuspended in 200 ul of NER buffer, vortexed for 15 sec and placed on ice with continued vortexing for 15 sec every 10 min for total 1 hour. Subsequently, the lysates were centrifuged at 17,000g for 10 min, and the supernatants (nuclear extract) were transferred into new tubes and an equal amount of ice-cold PBS and DNase I buffer (10x) was added. The extracts were split into two tubes - 20U DNase I (Thermo Fisher, 18068015) was added into one tube only. Reactions were incubated overnight at 37C. DNA was extracted following QIAamp DNA Mini kit (Qiagen) and eluted in 50uL water. qPCR was performed with 2uL of eluted DNA and the same parameters as described above in Vector copy number qPCR section. 1e9 vector copies of purified AAV treated in the same manner as the nuclei were used as control.

### Southern blotting

Nuclear gDNA was extracted from liver tissue as indicated above. Then gDNA was digested overnight with XhoI (NEB) that does not cut in the vector, but in the host gDNA. Digested DNA was run in a 1% TAE agarose gel at room temperature overnight. After electrophoresis, the gel was washed with denaturing buffer (3⍰M NaCl and 400⍰mM NaOH) twice for 5⍰min, and DNA was transferred to an Amersham Hybond-XL membrane (GE Healthcare) using transfer buffer (3⍰M NaCl and 8⍰mM NaOH) overnight. Membranes were washed with 2× saline sodium citrate (SSC) buffer for 5⍰min and blocked with UltraPure Salmon Sperm DNA (Thermo Fisher) in QuikHyb Hybridization Solution (Agilent Technologies) for 1⍰h at 65⍰°C. Probes for FLuc (500 bp) were generated using gel-purified PCR amplicons containing GFP sequence and a BcaBEST Labeling kit (Takara) and [α-32P]-dCTP (PerkinElmer), and probe hybridization were performed overnight at 65⍰°C with rotation. The membrane was washed with 2× SSC buffer and with 2× SSC containing 0.1% SDS at 65⍰°C. Signals were visualized using a Personal Molecular Imager System (Bio-Rad) and analyzed with Quantity One 1-D software (Bio-Rad).

### Flow cytometry

Cultured cells transduced with rAAVs were collected by trypsinization, washed with PBS and resuspended in cold PBS. Cells were kept on ice and protected from light until analyzed. Singlet cells were determined based on forward scatter/side scatter (FSC/SSC) plot, and GFP+ or TdTomato+ fractions were gated based on non-transduced cells, as negative control. Data was collected for 10’000 gated cells. The percentage of GFP+ or TdTomato+ expressing cells was evaluated using BD LSRII flow cytometer with BD FACSDiva software, and data were analyzed using the FlowJo software package.

### Cut&Tag

Reagents and protocol are commercially available by Epicypher (https://www.epicypher.com/products/epigenetics-reagents-and-assays/cutana-cut-and-tag-assays). See Supplementary Table 2, for details about reagents and antibodies used. Briefly, cultured cells transduced with rAAVs were collected by trypsinization and washed with PBS. Nuclei were extracted with NE buffer and mixed with activated Concanavalin A beads. After successive incubations with primary antibody (overnight) and secondary antibody (for 30 min) in antibody and Digitonin150 buffer respectively, the beads were washed with Digitonin150 buffer and resuspended in Digitonin300 buffer with 2.5uL of pA(G)-Tn5 for 1 hr and washed with the same buffer. Incubations were performed at room temperature in low-retention PCR strip tubes (Epicypher). Tagmentation was performed for 1 hr at 37C in Tagmentation buffer that provides MgCl_2_. Beads were washed with TAPS buffer and DNA material was released by adding SDS Release Buffer and incubating at 58C for 1hr. Quenching of SDS was performed by adding SDS Quench buffer and PCR was performed directly on this material. Universal P5 and indexed P7 primer solutions were used (see Supplemental Table 3 for sequences), and 21 cycles of PCR were performed. Clean-up was performed with AMPure beads, eluted in 15uL 0.1x TE buffer. Qubit and Bioanalyzer were used to verify library qualities before pooling samples for sequencing. The barcoded libraries were mixed to achieve equimolar representation aiming for a 10nM final concentration. Sequencing was performed with Illumina HiSeq 4000, 150cycles total per lane, 2×75 paired-end reads, depth of >2M reads per sample.

For in vivo Cut&Tag, snap-frozen liver tissue (~100⍰mg) was homogenized in RINO 1.5-ml Screw-Cap tubes filled with stainless steel beads and 1mL of NE buffer using a bead homogenizer (Next Advance Bullet Blender Storm - BBY24M) at speed 8 for 4 minutes. The homogenate was transferred into to a 50mL tube with 12mL of NE buffer and incubated on ice for 10 min. Nuclei were pelleted by a 5 min centrifugation at 600g, the supernatant was decanted, and the pellet was resuspended in 1mL of NE buffer. Nuclei were quantitated after Trypan Blue staining using an automated cell counter (Countess, Thermo Fisher) and diluted accordingly. 1e5 nuclei per target protein were used for Cut&Tag. The subsequent steps were identical to the protocol described above using nuclei from cultured cells.

### Tn5 normalizer - Illumina DNA prep

The Cut&Tag coverage plots of gAAV were normalized by total DNA coverage. Tn5 DNA libraries were created for all conditions (Extended Data Fig 2 c, 3 d), similar to what an input sample would be for chromatin immunoprecipitation. The mean coverage value of Tn5 libraries was used to divide the Cut&Tag coverage of all targets. This normalization is important, since there was a significant difference in the gAAV reads obtained in human vs mouse cells, independent of capsid.

To do this, cultured cells transduced with rAAVs were collected by trypsinization and washed with PBS. Nuclei were extracted for at least 5e5 cells with NE buffer following Epicypher Cut&Tag protocol. Total gDNA was extracted using a QIAamp DNA Mini kit (Qiagen) including RNAse treatment. The protocol for Illumina DNA prep was followed as per manufacturer’s instructions with 100ng of DNA as input, 30min tagmentation and 8 cycles of PCR. The same indexed primers as for Cut&Tag were used for PCR amplification. Equal volumes of each barcoded library (each around 10nM assessed by Bioanalyzer) were pooled and library quality and quantity was assessed using the Qubit and Bioanalyzer. Sequencing was performed with Illumina Novaseq SP100, 150cycles total per lane (2 lanes total), 2×75 paired-end reads, depth of >15M reads per sample.

### Data processing and analysis

For all figures with barplots, data was processed in GraphPad Prism 9, error bars represent standard error of the mean, and n = 3 or more biological replicas, except for Fig 1 e and Extended Data Fig 1 c. Figure 4 b-d and Extended Data Fig 5 c represent the ratios of means of at least 3 biological replicas.

For Cut&Tag a total of 42 (cells in culture) and 21 (liver tissue) different conditions with at least three replicas were processed and analyzed.

Reads were trimmed with Trimmomatic-0.39 program for classical Illumina adapters. Aligned using Bowtie2 version 2.2.5 (https://sourceforge.net/projects/bowtie-bio/files/bowtie2/2.2.5/) with options: --local --very-sensitive-local --nounal --no-mixed --no-discordant --phred33 -I 10 -X 700; mapping was performed using the hg38 or mm10 build of the human and mouse genome respectively, merged with viral genome sequence (gAAV). The viral genome sequence is the same as plasmid pAAV-CAG-FLuc from ITR to ITR. PCR duplicates were eliminated with “MarkDuplicates” command of Picard version 2.23 (https://broadinstitute.github.io/picard/index.html).

Tracks were made as bedgraph files of normalized counts, using deeptools 3.3 (https://deeptools.readthedocs.io/en/develop/content/list_of_tools.html) bamCoverage with options: --binSize 1, --normalizeUsing CPM; track plots were made with R package Gviz 4.1. Coverage was calculated as SUM[(normalized count*(end-start position+1))/gAAV size] for each dataset of Cut&Tag and Tn5 normalizer. Importantly, ITR regions were removed from the calculation, as their coverage is overrepresented in all samples, perhaps due to recognition of ITRs from ssDNA. For in vitro datasets, coverage was normalized as: gAAV coverage C&T/ Tn5 coverage of corresponding condition. Coverage boxplots with or without normalization were generated with R package ggplot2 4.1.2. Boxplots display the median (thick bar), two hinges (lower and upper hinges correspond to the first and third quartiles) and two whiskers. The upper whisker extends from the hinge to the largest value no further than 1.5 *IQR (inter-quantile range) from the hinge. The lower whisker extends from the hinge to the smallest value at most 1.5 * IQR of the hinge. Data beyond the end of the whiskers are called “outlying” points and are plotted individually.

Statistical analysis was performed by pair-wise comparisons of coverage means per condition and apply a t-test, using R package ggpubr 4.1.2.

## Data availability

All data needed to evaluate the conclusions in the study are present in the main text or the supplementary materials. Request for reagents should be directed to M.A.Kay. Processed data have been deposited in the Gene Expression Omnibus and are available under the accession number XXXXX.

---

**Extended Data Fig 1:**
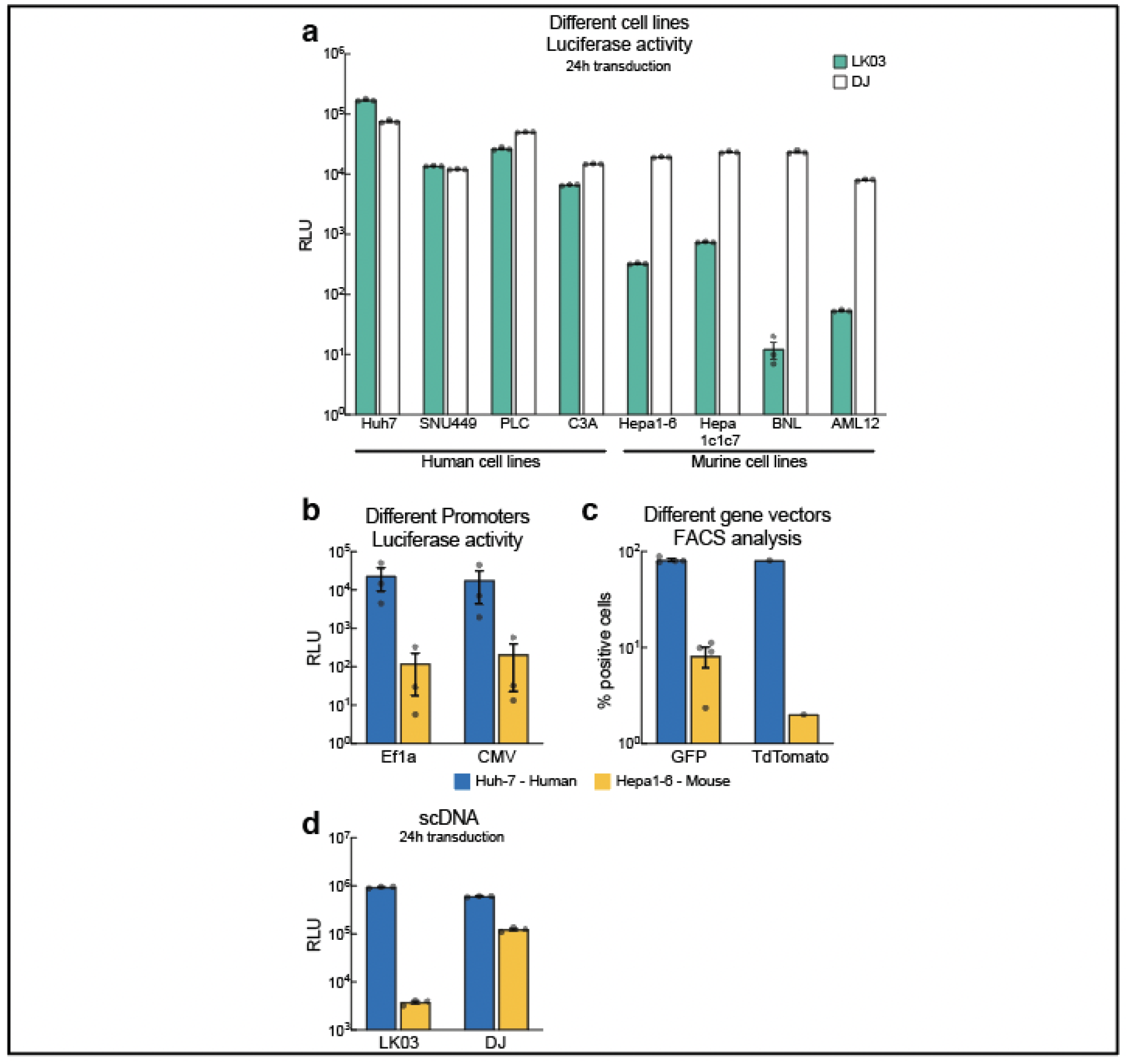
Transduction inefficiency of AAV-LK03 in murine cells is independent of cell line, promoter, transgene, or genome structure (scDNA). **(a)** Luciferase activity assay in various human and murine cell lines 24h post transduction using rAAV expressing FLuc cassette. **(b)** Luciferase activity in indicated cell lines 48h post transduction with AAV-LK03 packaged luciferase vectors with two different promoters (Ef1a or CMV). **(c)** FACS analysis of two cell line 48h post transduction with AAV-LK03 expressing GFP or TdTomato under control of the CAG promoter.**(d)** Luciferase activity assay of cells 24h post transduction with indicated capsids delivering a self-complementary (sc) RLuc vector.

**Extended Data Fig 2:**
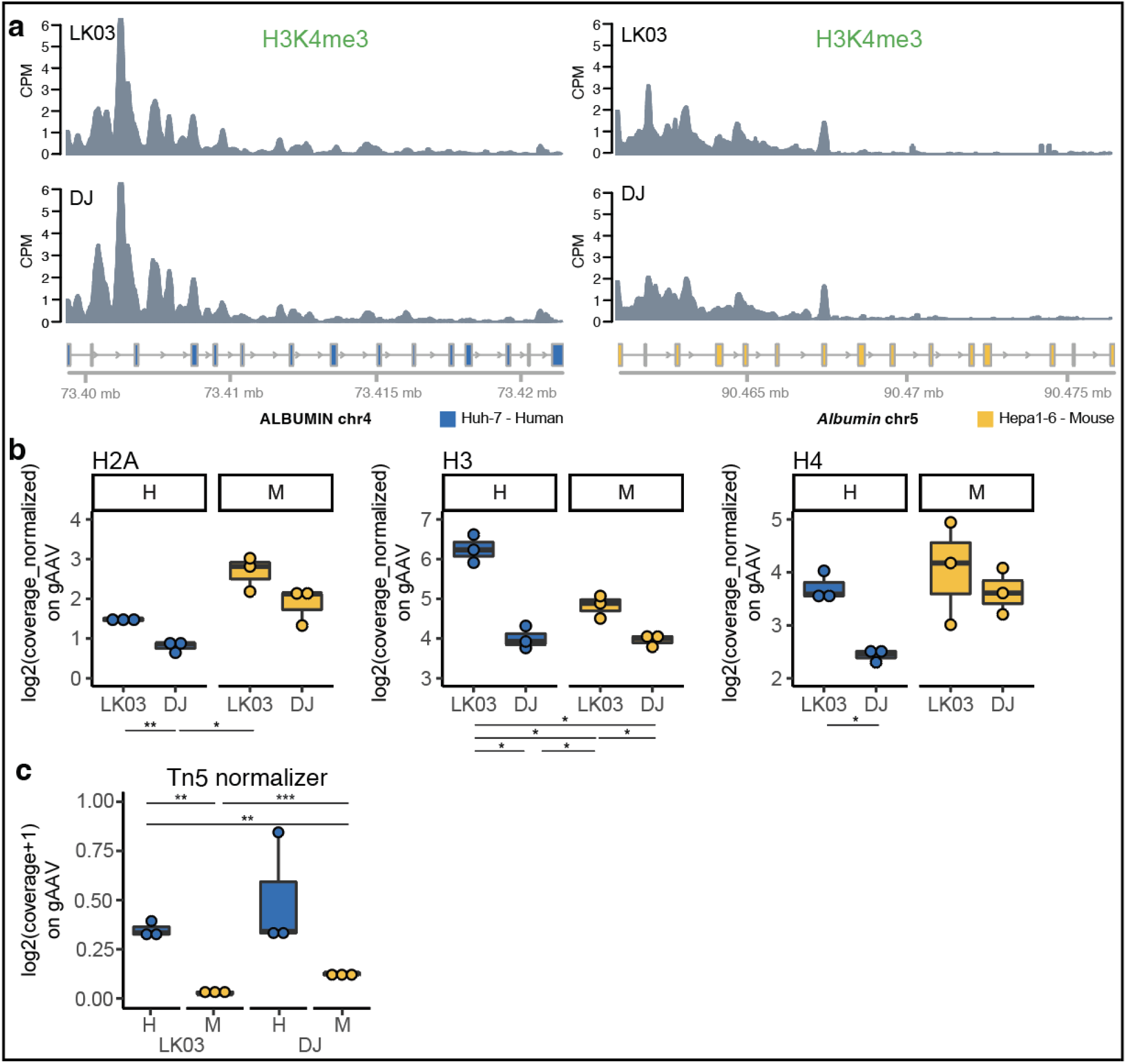
Host genome H3K4me3 signals are comparable and independent of AAV capsid while core histones enrichment in gAAV is variable between capsids and species. **(a)** H3K4me3 normalized (CPM) signal on the *Albumin* gene (as representation of host genome) obtained from the Cut&Tag assay and next generation sequencing for Huh7 human and Hepa 1-6 mouse cell lines transduced with AAV-LK03 and AAV-DJ. **(b)** Cut&Tag boxplots of normalized coverage on the AAV genome, for core histones in Huh7 human and Hepa 1-6 mouse cell line transduced with AAV-LK03 and AAV-DJ. **(c)** Tn5 Illumina DNA sequencing used as a normalizer value for Cut&Tag coverage for AAV-LK03 and AAV-DJ delivered gAAV. Statistic p-value * <0.05, ** <0.01, *** <0.001. H=human, M=mouse.

**Extended Data Fig 3:**
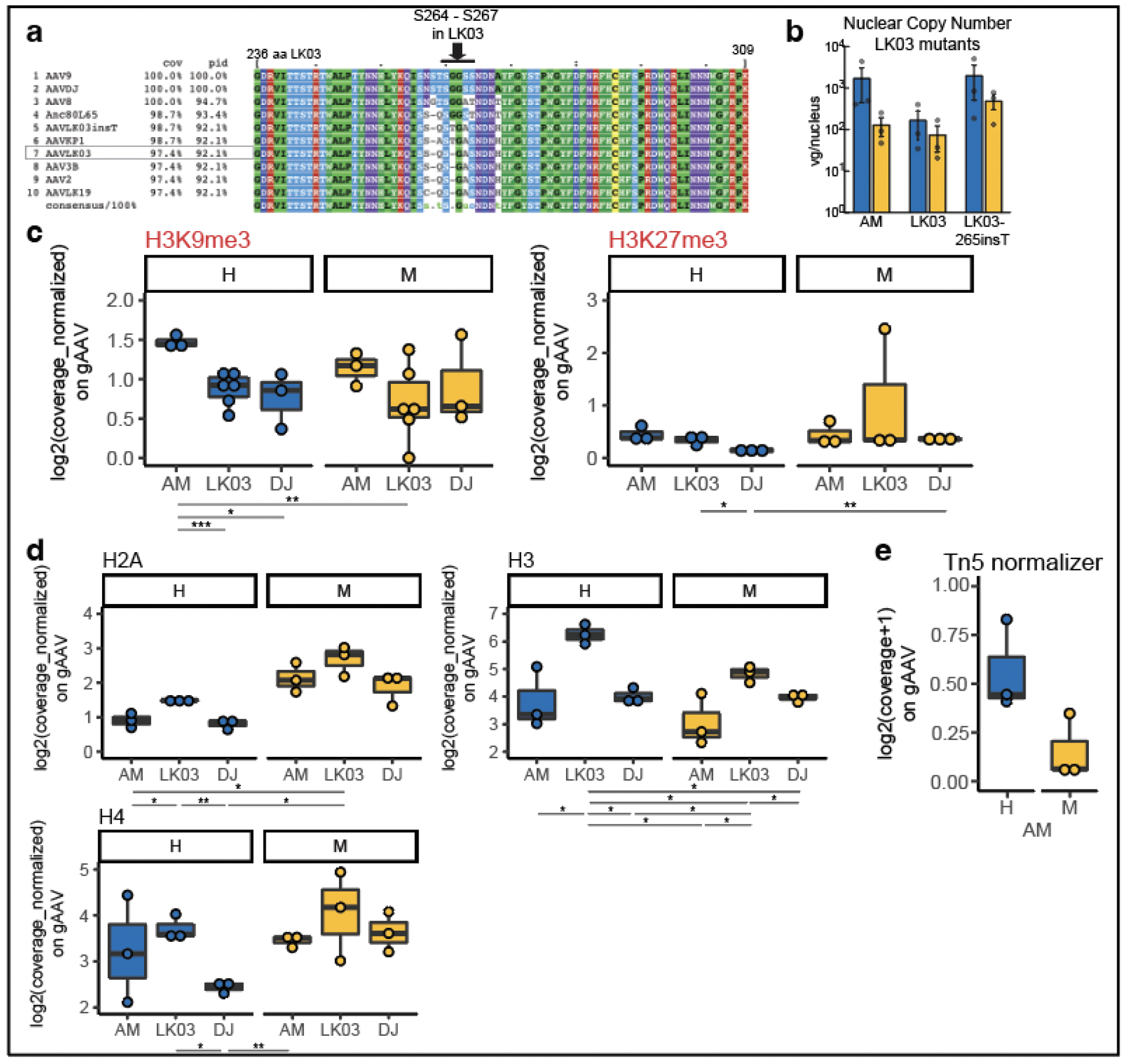
AAV-AM is a capsid with an insertion of a glycine amino acid in the AAV-LK03 capsid sequence. Inactive-related histone modifications are similarly enriched, and core histones have variable enrichments, on genomes delivered by AAV-AM as the other capsids. **(a)** Alignment of AAV-LK03 with multiple other capsids from amino acid 236-309, arrow points at variable region where Glycine was inserted at position 266 to create capsid AAV-AM. **(b)** Nuclear copy number qPCR quantification in Huh7 human and Hepa 1-6 mouse cells transduced with AAV-AM, AAV-LK03 and AAV-LK03-265insT. **(c)** Cut&Tag boxplots of normalized coverage on AAV genome, for inactive-related histone modifications (H3K9me3, H3K27me3) and **(d)** core histones (H2A, H3, H4) for Huh7 human and Hepa 1-6 mouse cells transduced with AAV-AM, AAV-LK03 and AAV-DJ for 48h. **(e)** Tn5 Illumina DNA sequencing, to use as normalizer value for Cut&Tag coverage for AAV-AM delivered gAAV. Statistic p-value * <0.05, ** <0.01, *** <0.001. H=human, M=mouse.

**Extended Data Fig 4:**
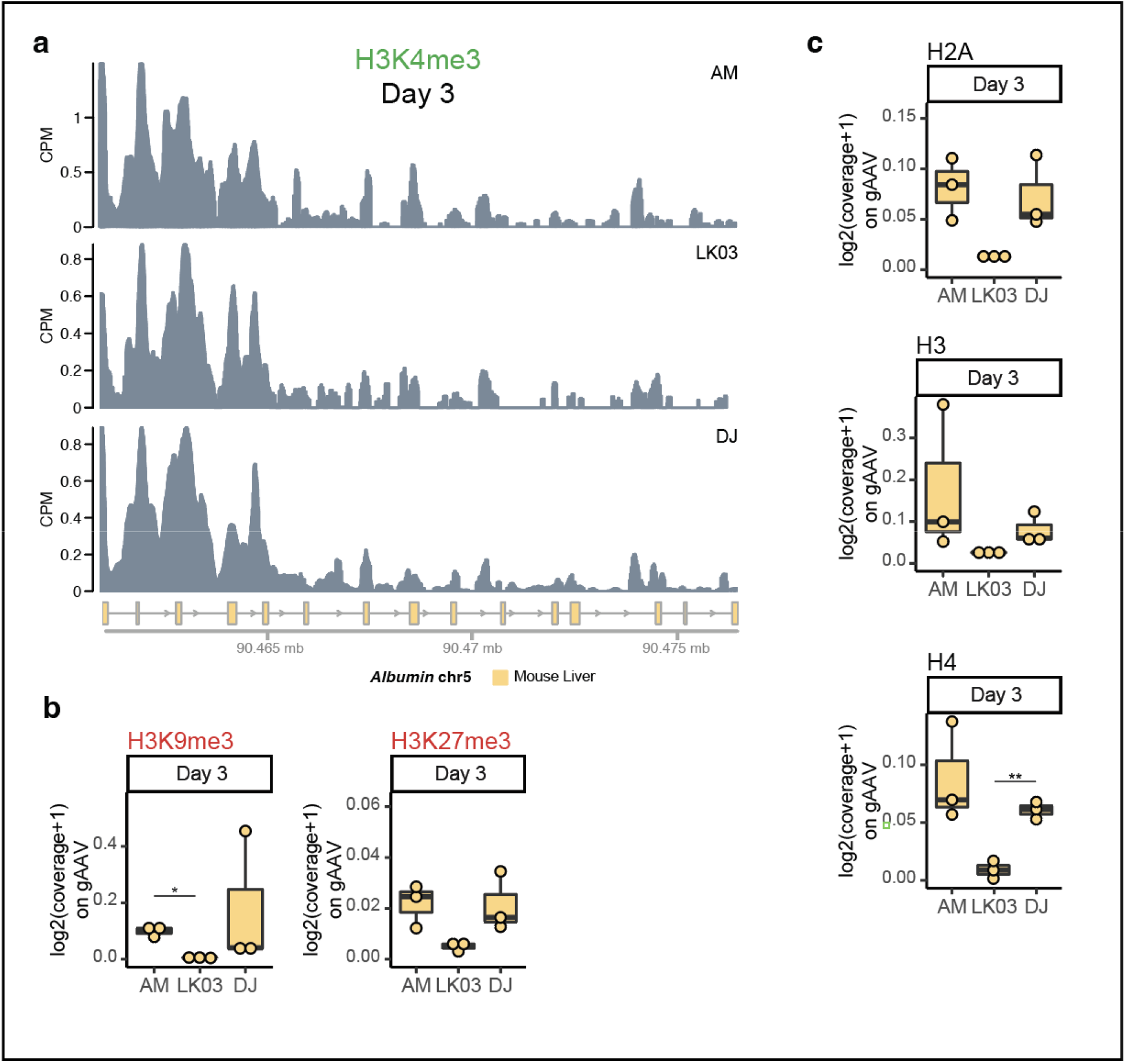
Host genome H3K4me3 signals are comparable independent of AAV transduction in vivo. Inactive-related histone marks and core histones enrichments are comparable between AAV-AM and AAV-DJ but not AAV-LK03 delivered genomes, in vivo. **(a)** H3K4me3 normalized (CPM) signal on the Albumin gene (as representation of the host genome) obtained from Cut&Tag assay and next generation sequencing for mouse liver transduced with AAV-AM, AAV-LK03 and AAV-DJ, collected 3 days post injection. **(b)** Cut&Tag boxplots of normalized coverage on AAV genome, for inactive-related histone modifications (H3K9me3, H3K27me3) and **(c)** core histones (H2A, H3, H4) for mouse liver transduced with AAV-AM, AAV-LK03 and AAV-DJ, collected 3 days post injection. Statistic p-value * <0.05, ** <0.01.

**Extended Data Fig 5:**
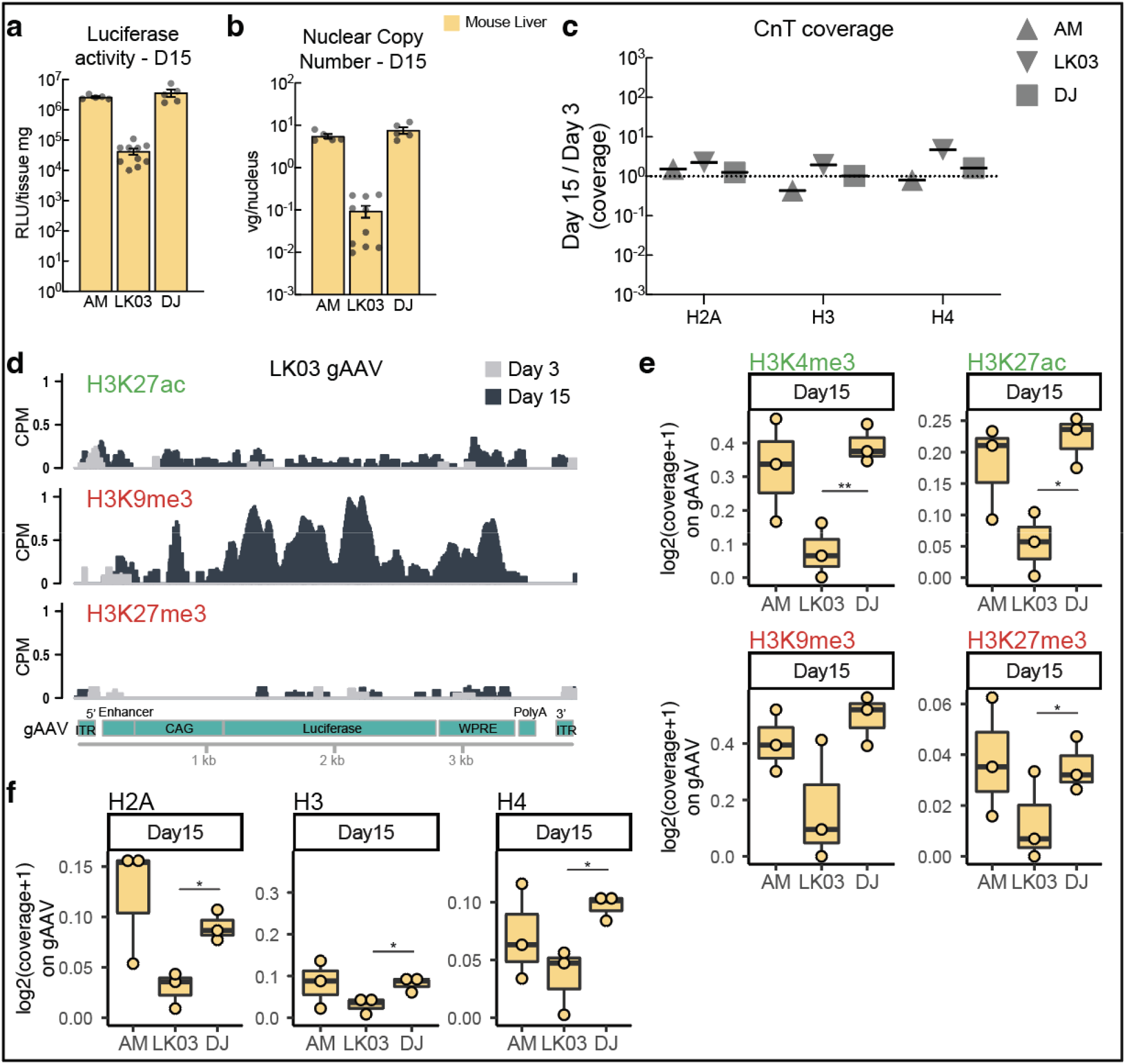
In vivo assays on mouse liver tissue at 15 days post injection. **(a)** Luciferase activity, **(b)** nuclear copy number qPCR of AAV genomes as indicated, collected 15 days post injection. **(c)** Ratio Day 15 vs Day 3 post injection of Cut&Tag coverage for core histones. **(d)** Comparison of Cut&Tag enrichment of indicated histone modifications on AAV-LK03 derived AAV genome at different days post injection. Normalization as CPM. **(e)** Cut&Tag boxplots of normalized coverage on AAV genome of histone marks and **(f)** core histones, collected 15 days post injection. Statistic p-value * <0.05, ** <0.01.

